# Binning meets taxonomy: TaxVAMB improves metagenome binning using bi-modal variational autoencoder

**DOI:** 10.1101/2024.10.25.620172

**Authors:** Svetlana Kutuzova, Pau Piera, Knud Nor Nielsen, Nikoline S. Olsen, Leise Riber, Alex Gobbi, Laura Milena Forero-Junco, Peter Erdmann Dougherty, Jesper Cairo Westergaard, Svend Christensen, Lars Hestbjerg Hansen, Mads Nielsen, Jakob Nybo Nissen, Simon Rasmussen

## Abstract

A common procedure for studying the microbiome is binning the sequenced contigs into metagenome-assembled genomes. Currently, unsupervised and self-supervised deep learning based methods using co-abundance and sequence based motifs such as tetranucleotide frequencies are state-of-the-art for metagenome binning. Taxonomic labels derived from alignment based classification have not been widely used. Here, we propose TaxVAMB, a metagenome binning tool based on semi-supervised bi-modal variational autoencoders, combining tetranucleotide frequencies and contig co-abundances with contig annotations returned by any taxonomic classifier on any taxonomic rank. TaxVAMB outperforms all other binners on CAMI2 human microbiome datasets, returning on average 40% more near-complete assemblies than the next best binner. On real long-read datasets TaxVAMB recovers on average 13% more near-complete bins and 14% more species. When used in a single-sample setup, TaxVAMB on average returns 83% more high quality bins than VAMB. TaxVAMB bins incomplete genomes drastically better than any other tool, returning 255% more high quality bins of incomplete genomes than the next best binner. Our method has immediate research and industrial applications, as well as methodological novelty which can be translated to other biological problems with semi-supervised multimodal datasets.

---

Shotgun metagenome sequencing is an accessible technology that enables high-throughput analysis of complex microbial communities for both taxonomic profiling and metagenome assembly tasks. The field is currently dominated by short-read (commonly 100–300 bp) technologies ^1^, however, recently long-read sequencing has gained prominence, as it allows the recovery of even more individual genomes with higher accuracy ^2–4^. When working with environmental samples in the absence of cultured isolates, the assembled contigs are grouped together during the process of metagenome binning ^5^.

Most metagenome binning tools ^6–11^ are based on analysing both contig composition, commonly represented as k-mer frequencies vectors such as tetranucleotide frequencies (TNFs) ^12^, and contig co-abundances across multiple samples. Besides the information contained in contigs, some tools additionally rely on assembly graphs ^13–16^, codon usage ^17^, GC content ^6^, single-copy genes ^18–22^ and contig-level taxonomy profiling ^20,23–25^. Furthermore, ensemble tools leverage the binning results created by multiple approaches ^26–28^. Most metagenome binning tools have been optimized for short-read sequences and their performance on long-read datasets has not been thoroughly evaluated. Recently, several tools such as GraphMB ^15**?**^, SemiBin2 ^21^ and LRBinner ^29^ have been developed specifically for long-read sequencing data. In general, the large amounts and complexity of metagenomic data make it a suitable application for deep learning algorithms ^11,15,16,20–22,28^.

For the purpose of this study, we emphasise a rough distinction between the intrinsic features ^30^ derived purely from a given set of reads and their corresponding contigs (k-mer frequencies, GC-content, co-abundances) and the annotation features that require searching external databases (e.g. single-copy genes, taxonomic labels from sequence alignment). A taxonomic label is an example of an annotation feature, which can be extracted from a read or a contig using taxonomic profiling tools ^31–38^. These annotations are often incomplete since not all the contigs can be successfully mapped to a reference sequence. The annotation might also be biased towards better-studied organisms that will be more prevalent in databases.

Recently, single-copy genes (SCGs) have been used as a key clustering feature by the SemiBin2 ^21^ and Comebin ^22^ methods. Traditionally, SCGs have been used for evaluating metagenome assembled genomes (MAGs), as in the popular metagenomic binning evaluation tools CheckM and CheckM2 ^39,40^. While missing or duplicated single-copy genes are indeed a strong signal of the MAG quality, one might be cautious about using these both as input to binning and as an evaluation of the produced bins. This turns the evaluation metric into a training target, an observation sometimes referred to as Goodhart’s law: “When a measure becomes a target, it ceases to be a good measure” ^41^.

For instance, SemiBin2 performs self-supervised deep learning using only intrinsic features (TNFs and co-abundances) followed by a reclustering step using SCGs as the optimization target ^21^. The reclustering step includes a greedy search over multiple possible clusterings, selecting clustering with the best contamination and completeness values calculated based on the presence of single-copy genes. In principle, reclustering does not have to be done on the embeddings produced by a deep learning model, but could more efficiently be applied directly to the input features, foregoing the embedding entirely. The authors of SemiBin2 reported that the performance of reclustering the input features of simulated short-read datasets was only 17.7% lower than that of reclustered embeddings. This suggests that reclustering, which introduces the evaluation metrics as an input to the model, is by far the most informative part of the SemiBin2 algorithm. Therefore, investigation of binning performance with and without single-copy genes is needed to provide unbiased benchmarks of different methods.

Incorporation of taxonomic information presents a computational challenge due to its hierarchical nature. The taxonomic labels used to classify the hierarchical phylogeny of microorganisms are organised into the seven classical taxonomic ranks from kingdom to species. Lower taxonomic ranks provide more precise information about the contig phylogenetic placement, but are more often mislabeled or missing. As demonstrated in the Taxometer tool ^42^, a hierarchical loss allows training on the labels acquired on all the taxonomic ranks (e.g. phylum or genus) without requiring annotations on a particular taxonomic rank (e.g. species). Previously, both SemiBin ^20^ and SolidBin ^25^ used taxonomic labels to generate cannot-link constraints in the loss functions of self-supervised deep learning algorithms but neither labels themselves nor their hierarchical structure were a part of the training data.

A key feature of semi-supervised machine learning is that the models can be trained using both annotated and un-annotated samples. Analogous to the standard unsupervised VAEs, the semi-supervised multi-modal VAEs exhibit generative capabilities and produce embeddings for downstream tasks that combine the information from two or more modalities ^43–51^. Therefore, unlike other popular DL based methods with multimodal capabilities like stacked autoencoders or siamese networks ^52^, most multi-modal VAEs do not require the dataset to be fully labeled.

Here, we introduce TaxVAMB, which combines the strengths of intrinsic and annotation features to create high-quality MAGs that cover more taxonomic diversity than any other binning tool (Figure 1). It outperforms all other binners in the number of near-complete assemblies for both short- and long-read datasets. We demonstrate that using TaxVAMB is especially beneficial for datasets with fewer than 100 samples, and that it bins incomplete genomes drastically better than any other tool. TaxVAMB also runs sufficiently fast to be one of the few binning tools that can process large-scale experiments with as many as 1000 samples. We demonstrate the model performance on several short- and long-read datasets from various environments and found that TaxVAMB was 40.2% and 25% better compared to existing methods. TaxVAMB and the source code is freely available at https://github.com/RasmussenLab/vamb

**Fig. 1.**
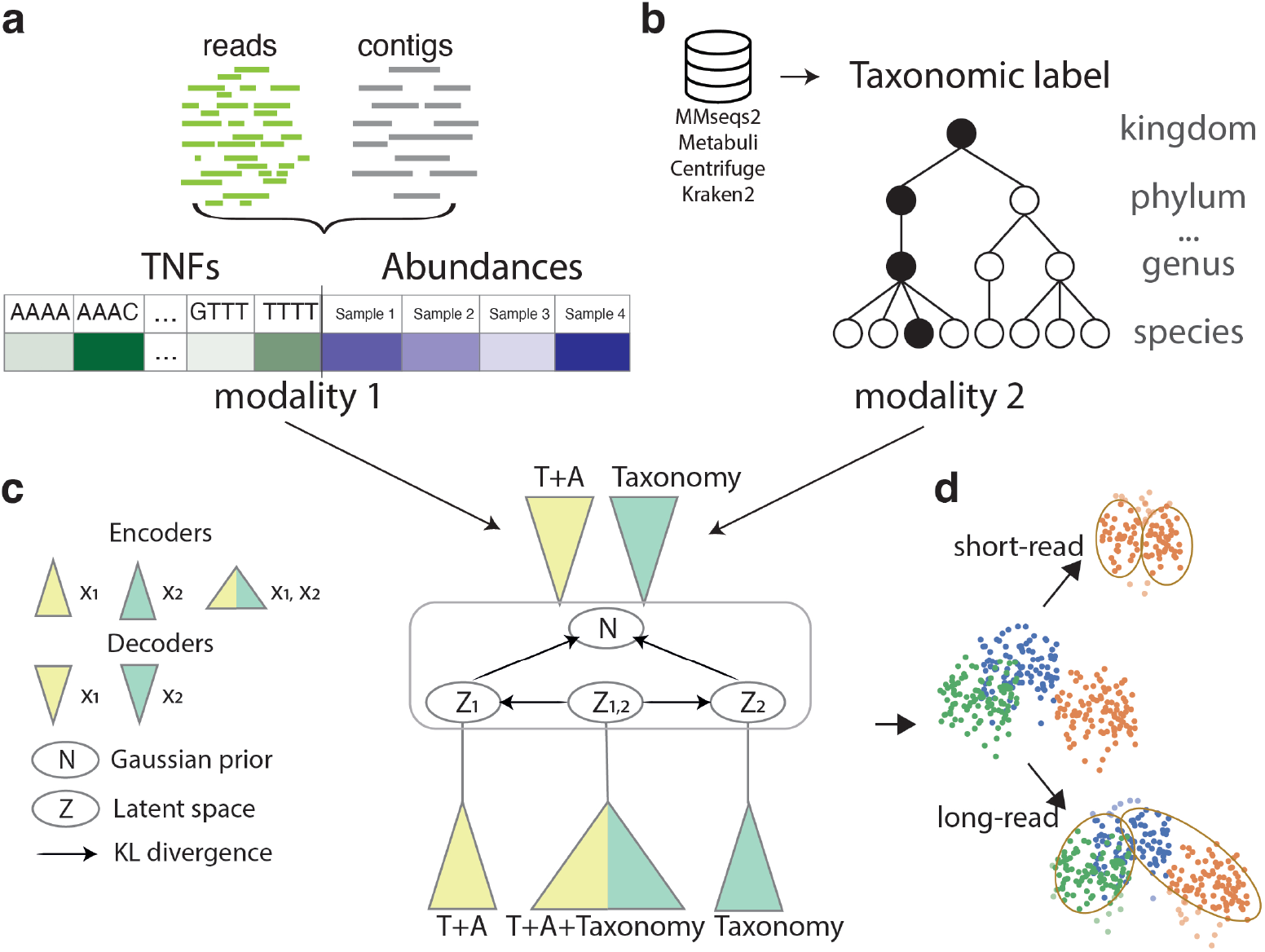
TaxVAMB workflow. **a**. Tetranucleotide frequencies and contig abundances across samples are extracted from reads and their assemblies. **b**. Contigs are annotated with taxonomic labels by a taxonomic classifier, and the labels are refined by the Taxometer tool, resulting in higher quality annotations. The taxonomic label is represented by a binary vector where each element encodes a taxon. Taxa from all the taxonomic ranks are present. **c**. We consider a concatenated vector of TNFs and abundances to be the first modality and the taxonomy label to be the second modality. Bi-modal variational autoencoder is trained on the two modalities. For each sample, three observations are created: 1) modality one; 2) modality two; 3) a concatenation of modality one and modality two. Each observation is encoded with a corresponding encoder. Each modality is decoded with its own decoder. The loss function has KL-divergence terms to ensure the shared representation regardless of the modality. See Methods for details. **d**. After training, clustering is performed on the resulting embedded vectors. The clustering method is based on iterative clustering as is used in VAMB. Optionally, a reclustering step using single-copy genes can be applied. Here, k-means based reclustering is used when the input is short-read data, and DBSCAN based reclustering is used if the input is long-read data.

## Results

TaxVAMB is a semi-supervised deep learning method that consists of two neural networks. First, we bridge intrinsic and annotation features by predicting taxonomy labels for unannotated contigs and refining the annotated ones using Taxometer trained on the contigs of this dataset ^42^. Second, we use labels on all the taxonomic ranks and the intrinsic features as two modalities to train a bi-modal variational autoencoder to construct the joint latent space. Bi-modal variational autoencoders are a family of VAE-based methods adapted for semi-supervised learning (Supplementary Figure S1). We use an architecture with three encoders (TNFs and abundances; taxonomic labels; TNFs, abundances and taxonomic labels) and two decoders (TNFs and abundances; taxonomic labels). The loss function is designed in a way to ensure that the similarity between the latent vectors produced by either of the three encoders is preserved if they are generated from the same contig. Following the strategy previously introduced in the Taxometer method ^42^, we use a flat softmax hierarchical loss ^53^ to train on all the taxonomic labels returned by a taxonomic classifier. The latent space is clustered using the original VAMB clustering algorithm. Optionally, the latent space can also be reclustered using the method adapted from SemiBin2, which is based on single-copy genes (Figure 1, Supplementary Figure S2, Supplementary Table 1).

### TaxVAMB outperformed all other binners on CAMI2 Toy datasets

To evaluate TaxVAMB’s performance on human microbiome datasets that include truth annotations, we benchmarked TaxVAMB against six other binners on the synthetic CAMI2 toy human microbiome short-read datasets. We used BinBencher v0.3.0 ^54^ to compute two distinct metrics relative to the known ground truth: The number of near-complete genomes, and the number of near-complete assemblies (see Methods). Measured in the number of near-complete assemblies, TaxVAMB outper-formed all datasets with improvement over the second best binner of 90% for Airways (272 over 138 from VAMB), 24% for Urogenital (158 over 127 from AVAMB), 9% for Gastrointestinal (178 over 163 from SemiBin2), 38% for Skin (272 over 184 from Comebin), and 40% for the Oral dataset (274 over 195 from VAMB) (Figure 2, Supplementary Figure S3, Supplementary Figure S4). Measured in the number of near-complete genomes, TaxVAMB demonstrated state-of-the-art performance on 4 out of 5 datasets. We found that the improvement of using TaxVAMB compared to the second best binner were 11% for Airways, 4% for Urogenital, 3% for Gastrointestinal, and 9% for Skin, whereas for the Oral dataset, the AVAMB binner was 3% better. The results showed the largest boost in performance metrics was when the recall was calculated using the contigs that were provided as an input to the binner, as opposed to the full genome. This indicates that TaxVAMB was drastically better at binning contigs that originated from incomplete genomes. For instance, TaxVAMB reconstructed 149 and 96 assemblies that had less than 0.9 of the total genome present in the input data for the Airways and Oral datasets, respectively. In comparison, SemiBin2 reconstructed 1 and 5 assemblies, respectively. For genomes that were almost completely present in the input data the other binners had nearly as good performance. We conclude that TaxVAMB reaches the state-of-the-art binning performance.

**Fig. 2.**
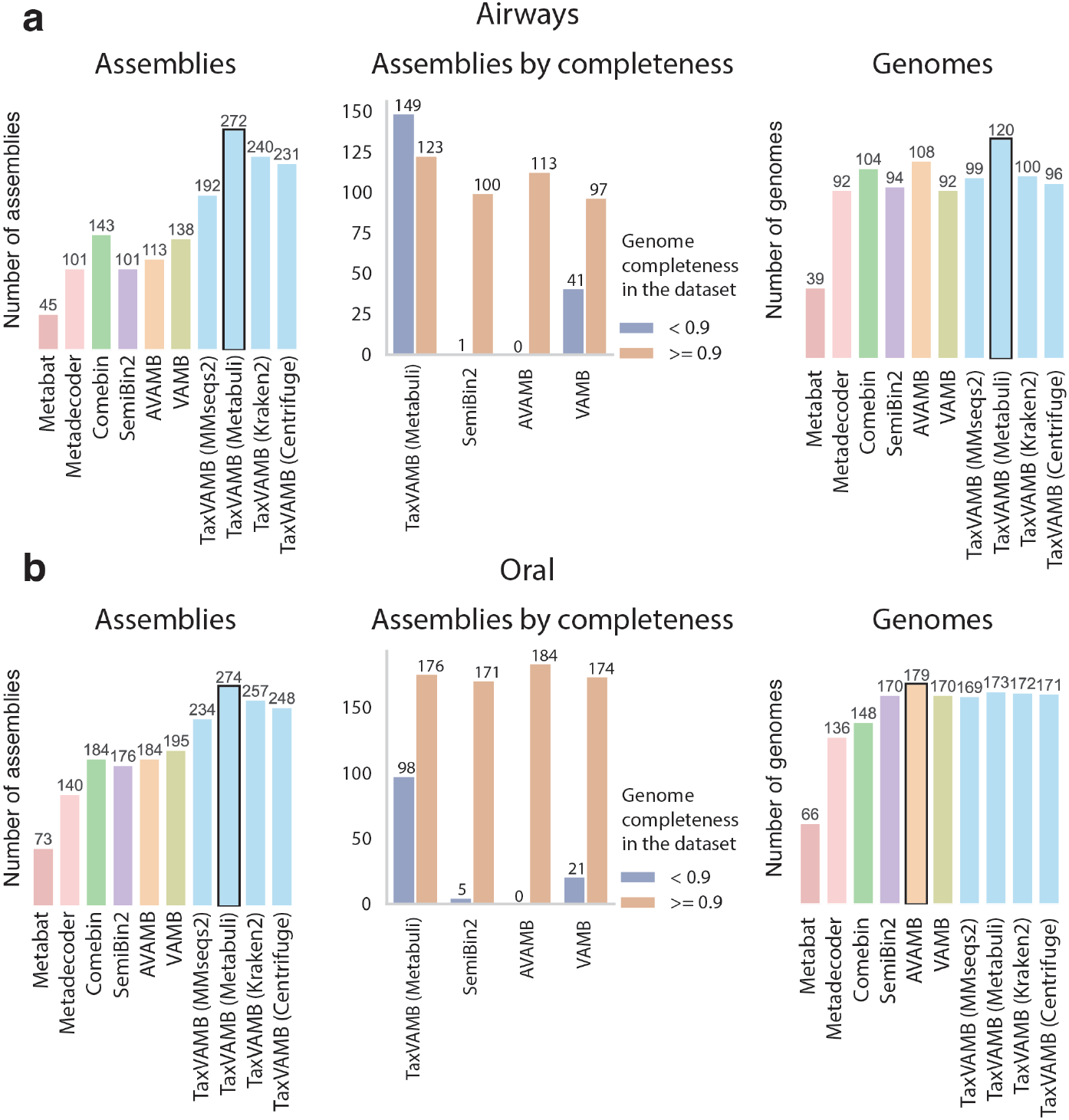
CAMI2 human microbiome benchmarks. Metagenome binning benchmarks on CAMI2 human microbiome datasets, with TaxVAMB using four different taxonomic classifiers. The bars show the number of near complete assemblies or genomes (recall more than 0.9 and precision more than 0.95). The ‘assembly’ and ‘genome’ metrics differ in how the recall is measured. The assembly metrics measures recall in respect to the part of the genome that was provided as input to the binner. The full genome metrics measures recall in respect to the full bacterial genomes even though parts of the genome can be missing from the assembly. Assemblies by completeness show the performance of the binners stratified by whether the genomes had an assembled share (contigs of that genome provided as input to the binner) less than 0.9, or equal/more than 0.9. SemiBin2, VAMB, AVAMB and TaxVAMB results are shown after applying the k-means based reclustering step. The datasets used are **a)** CAMI2 Airways; **b)** CAMI2 Oral.

### MAGs quality depends on the annotation tool

Because TaxVAMB used taxonomic annotations as input we evaluated how taxonomic annotations from different tools impacted the performance. Since the quality of the binning depended on noisy and incomplete taxonomic labels, the flexibility of Tax-VAMB is beneficial in case the input dataset is labeled inconsistently by different classifiers. In this case it is possible to run TaxVAMB independently for each taxonomic classifier and compare the quality of the resulting bins, even if the labels came from different databases such as GTDB or NCBI annotations. When investigating annotations provided by MMseqs2, Metabuli, Kraken2 and Centrifuge, we found that Metabuli resulted in the highest quality bins for 3 out of 5 the CAMI2 datasets providing 5%-20% more genomes. For the Gastrointestinal and Oral datasets the number of NC genomes differed by only 1 bin for Metabuli and MMseqs2 (Figure S4 - Gastrointestinal, Oral). Additionally, as Metabuli provided labels at the subspecies level, we included an additional benchmark where these annotations were used as bin identifiers. Here we found that TaxVAMB using Metabuli labels outperformed Metabuli itself with a 16% improvement on average for the five CAMI2 datasets measured in the number of NC genomes and with a 13% improvement on average measured in NC assemblies (Supplementary Figure S5).

### TaxVAMB outperformed SCG-based binners without using SCGs

Because single-copy genes are powerful features for binning we wanted to provide an unbiased estimate of performance across binners. To do this, we investigated the number of NC genomes reconstructed by VAMB, SemiBin2 and TaxVAMB before and after SCG-based reclustering. We observed that VAMB, which only uses intrinsic features, outperformed SemiBin2 before reclustering for all five datasets (Figure 3). Improvements of TaxVAMB compared to SemiBin2 were 28% for Airways, 11% for Urogenital, 1.7% for Oral, and 24% for Skin, whereas for the Gastrointestinal dataset, the SemiBin2 binner was 2.6% better. This suggests that a main factor of performance gain was from using single-copy genes for re-clustering of bins. This opposes the claim made in the SemiBin2 study that self-supervised contrastive learning using intrinsic features led to better MAGs. Conversely, we found that the performance of TaxVAMB performance was not that affected by reclustering using SCGs. Here we found that reclustering only resulted in 1.6%-4.1% more genomes when applied to TaxVAMB compared to 30%-110% more genomes when applied to SemiBin2. We conclude that, even when reclustering was applied, the performance of TaxVAMB was less driven by SCGs compared to SemiBin2.

**Fig. 3.**
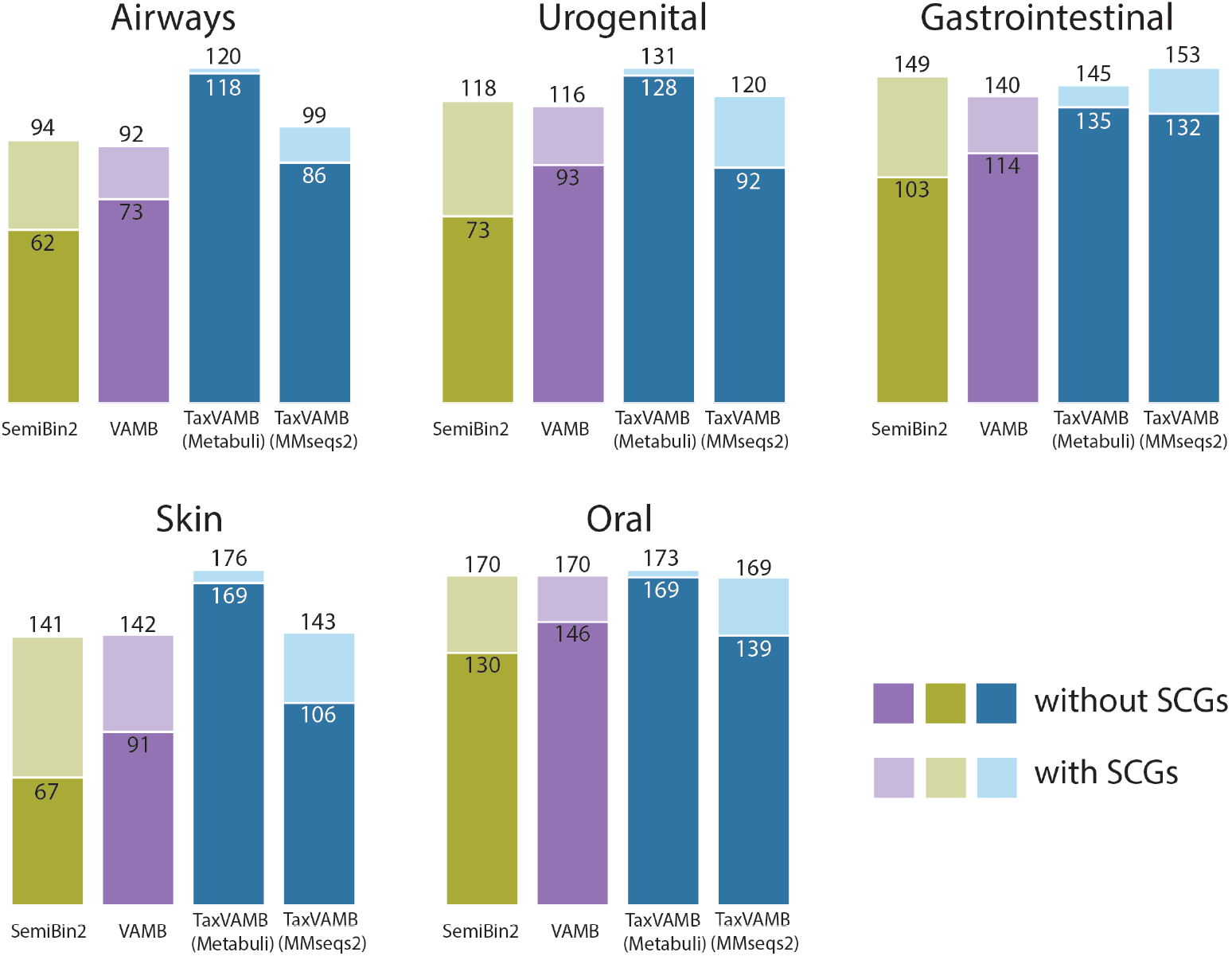
The effect of reclustering using single-copy genes. Evaluating the effect of reclustering using single-copy genes for SemiBin2, VAMB and TaxVAMB. The darker colors represent binning results without SCGs and the lighter colors represent the results using k-means based SCG-reclustering.

### TaxVAMB outperformed all other binners on a human gut long read dataset

To test the hypothesis that the performance gains of TaxVAMB would be higher for better-quality taxonomic labels, we benchmarked TaxVAMB on two long read datasets, contrasting a well-studied environment (human gut, three samples) with a poorly studied environment (sludge from an anaerobic digester, four samples). Given that the human gut is more well-studied compared to digested sludge, we hypothesized that the taxonomic classifiers would return more complete and correct labels for the human gut samples compared to the sludge samples. As expected, for the human gut dataset TaxVAMB outperformed SemiBin2 and AVAMB, reconstructing 25% more near-complete bins (Figure 4a). When applied to the sludge dataset, we found that TaxVAMB performed similarly to VAMB and AVAMB, reconstructing 248, 252 and 248 NC genomes, respectively. In comparison, SemiBin2 reconstructed 223 NC genomes from the sludge data. This confirms that a potentially noisy and incomplete taxonomy did not carry sufficient signal to improve or degrade binning performance of TaxVamb. As with the short-read datasets above, we evaluated the effect of reclustering using single-copy genes (see Methods subsection 6). For the sludge dataset, the application of SCG based reclustering improved performance for both VAMB and TaxVAMB. For the human gut dataset, TaxVAMB generated the most near-complete bins without using single-copy genes, and the performance was even reduced when using reclustering (Figure 4a). We then investigated the phylogenetic diversity of the NC bins reconstructed by SemiBin2, VAMB, and TaxVAMB from the human gut dataset ^55^. (Figure 4b). At phylum level TaxVAMB recovered 9 phyla, VAMB recovered 8 and SemiBin2 recovered 7. Additionally, TaxVAMB recovered 27% more species than SemiBin2 at the species level. Most of the MAGs of the sludge dataset were assigned to a novel species whereas novel species were rare in the human gut dataset (Figure 4c). The results suggested that TaxVAMB, using the high-quality taxonomy as input to binning, recovered more diverse MAGs.

**Fig. 4.**
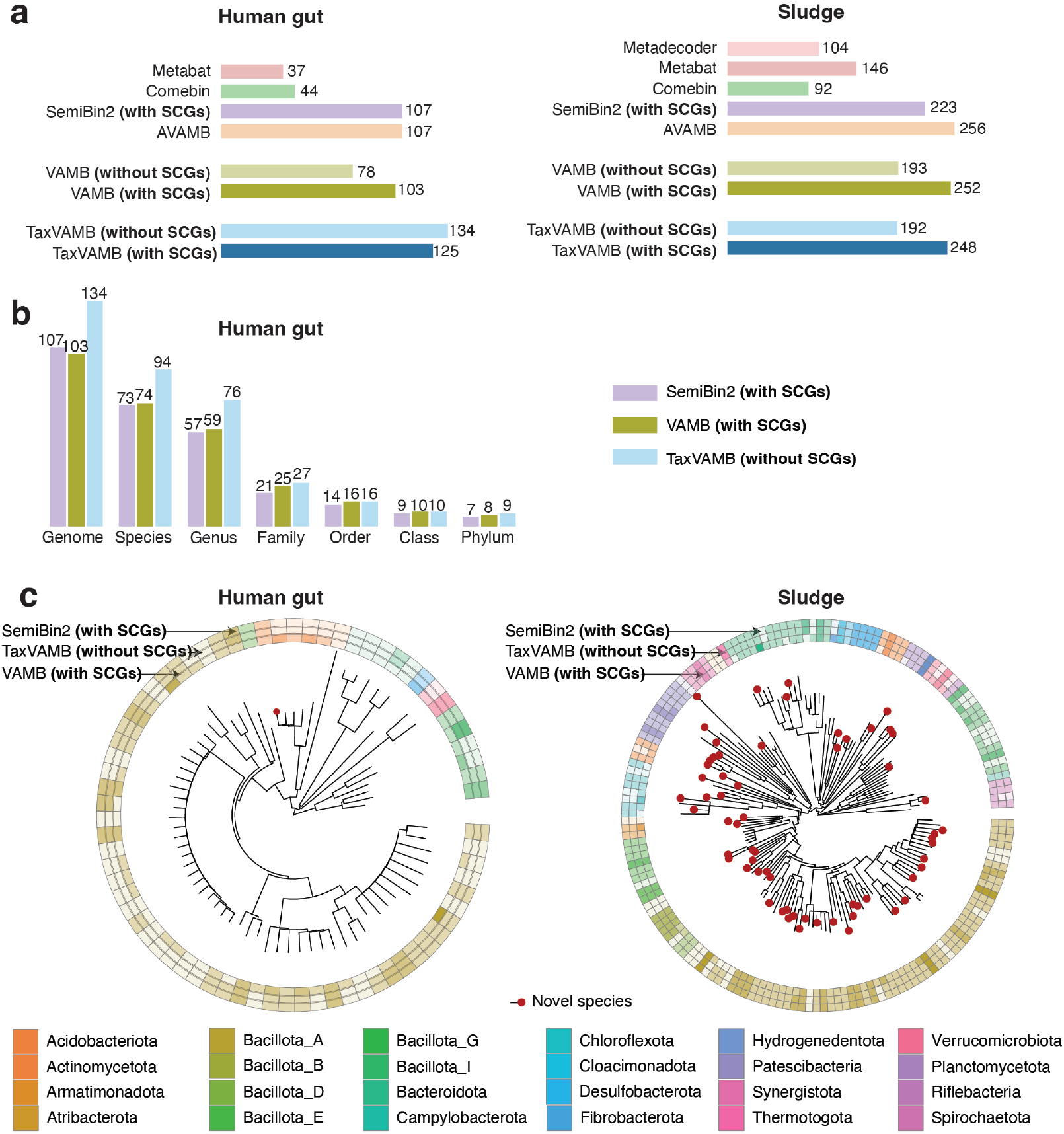
Long-read datasets benchmarks. **a**. Human gut and sludge dataset benchmarks for different metagenomic binners. The performance is measured as the number of ‘near-complete bins’, i.e. bins evaluated by CheckM2 to have > 90% completeness and < 5% contamination. VAMB and TaxVAMB are presented with and without SCG-based DBSCAN reclustering. **b**. The phylogenetic diversity of the SemiBin2, VAMB and TaxVAMB near-complete bins, using GTDB-tk placement, as the unique number of taxa on each taxonomic rank. SemiBin2 and VAMB were run using SCG-based DBSCAN reclustering, whereas TaxVAMB was run without. **c**. Visualisation of GTDB-tk placement for SemiBin2, VAMB and TaxVAMB down to the species level, annotated by the color on the phylum level. The darker color in the annotation indicates that more near-complete bins were recovered for this phylum. Red dot indicates a novel species.

### Bi-modal variational autoencoder outperformed stacked autoencoder in the number of high quality bins

To ensure that the semi-supervised architecture of the bi-modal VAE was beneficial for binning we empirically evaluated it compared to a stacked autoencoder (Figure S1). Using the short-read and long-read datasets presented above we found that TaxVAMB and the stacked VAE had similar performance with an average 2.3% absolute difference in performance across the short read datasets. The bi-modal VAE outperformed the stacked VAE 6 out of 15 times for MMseqs2, Kraken2 and Centrifuge classifiers. When Metabuli taxonomic labels were used, the bi-modal VAE always outperformed the stacked VAE with an average gain of 16% across all CAMI2 datasets (Supplementary Figure S6a). For the long-read datasets, TaxVAMB outperformed the stacked autoencoder architecture across all datasets and classifiers, with an average gain of 13% more high quality genomes for the Human Gut dataset and 18% more high quality genomes for the Sludge dataset (Supplementary Figure S6b). This indicate that binning the latent vectors provided by TaxVAMB using a bi-modal VAE resulted in better overall performance for the task of metagenomics binning compare to the same workflow that uses a stacked VAE.

### Taxonomy primarily improved binning at low number of samples

The benefit of using co-abundance for binning positively correlates with the number of samples in the experiment ^56^. We, therefore, investigated how taxonomic labels improved binning as a function of the size of the dataset. To achieve this, we varied the number of samples from 1-1000 when using human gut microbiome samples from Almeida et al. ^57^ comparing VAMB to TaxVAMB ((Figure 5a, see Methods for details). We found that when the dataset included 1,000 samples TaxVAMB only recovered 3% more near-complete MAGs than VAMB. However, when we subsampled the datasets to 100 samples, the performance improvement increased to 16%, while for datasets of 10 samples, the improvement was 23%. Finally, when performing single-sample binning, i.e. one sample as input, the performance increased by 48% when using TaxVAMB compared to VAMB (Figure 5b). In a similar experiment using a wheat phyllosphere dataset we found that the gains compared to single-sample binning were even larger, resulting in 118% more bins in TaxVAMB compared to VAMB. Interestingly, for the experiment using datasets of 10 samples, we found that TaxVAMB increased the number of near-complete bins to the level of running VAMB on chunks of 100 samples. Therefore, in experiments with few samples, TaxVAMB was able to compensate for a less expressive abundance vector by using the taxonomy label modality. We conclude that TaxVAMB delivered larger gains compared with VAMB on datasets with fewer than 100 samples.

**Fig. 5.**
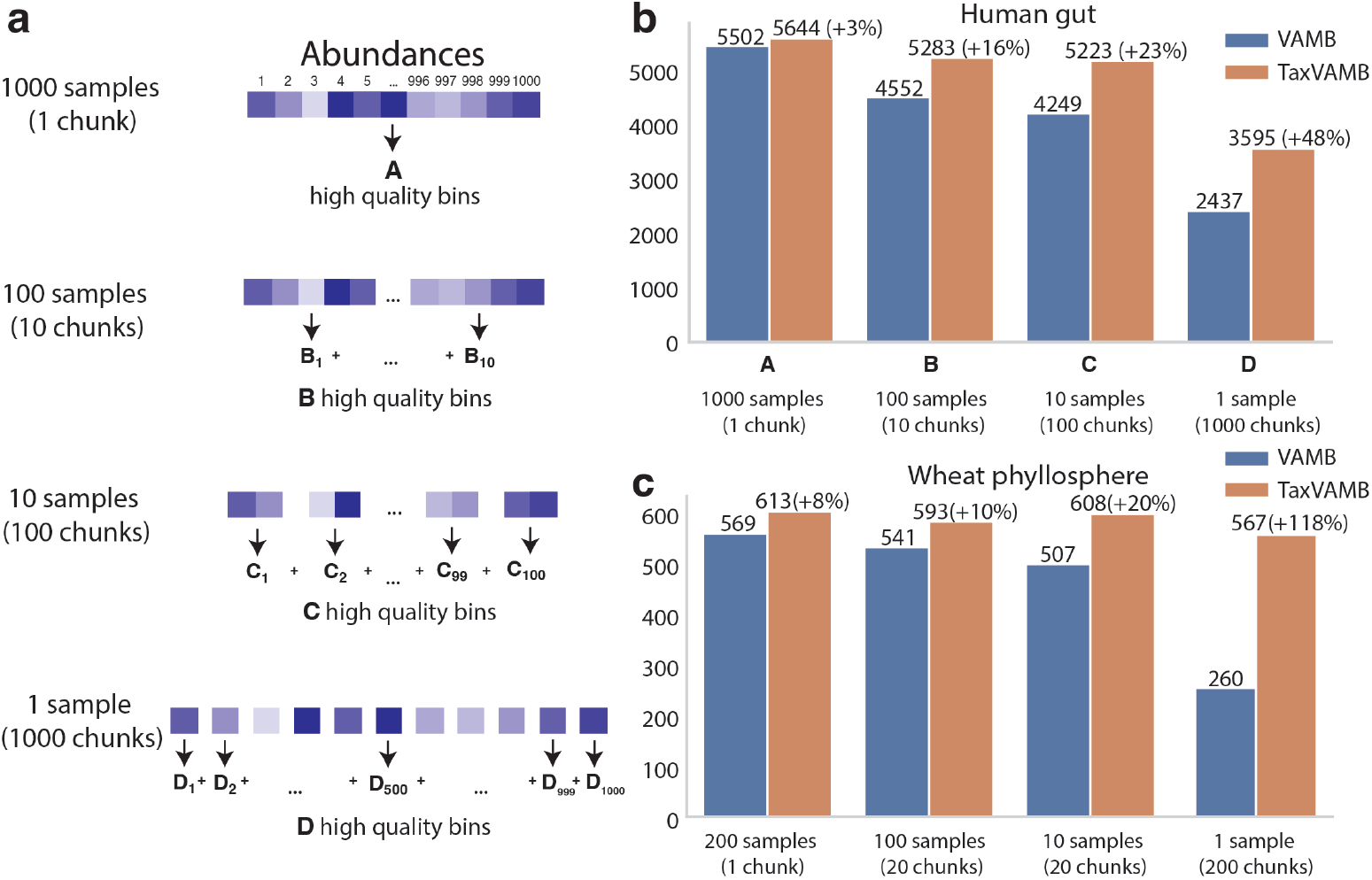
Effect of abundance vector and taxonomy information. **a**. For 1,000 samples, a single TaxVAMB and VAMB run was performed using all contigs and the entire abundance vector. 10 runs were performed on chunks of 100 samples and their corresponding contigs and abundances, 100 runs with 10 samples and 1,000 runs with 1 sample. The number of near-complete bins for all chunks was summed for each set of 1,000 samples. **b**. The results for the human gut dataset of Almeida et al. using TaxVAMB and VAMB. **c**. The results for the wheat phyllosphere dataset using TaxVAMB and VAMB.

### TaxVAMB provided consistent bin annotations

A key step of TaxVAMB is to predict taxonomic assignments of contigs without taxonomic labels. This is done using a neural network (Taxometer) resulting in taxonomic labels for all binned contigs (Supplementary Figure S7a). We, therefore, investigated if majority voting of these could be used as a taxonomic classification of the bins. Using Kraken2 classification of the CAMI2 human microbiome dataset we isolated the subset of high-quality MAGs. We found that 95-98% of bins per dataset were correctly annotated down to the species level, with the GTDBtk classifier ^55^ correctly annotating 98-99% bins (Supplementary Figure S7b, Supplementary Figure S8). TaxVAMB annotations have the advantage over GTDBtk of not requiring any additional runtime and not being limited by prokaryotes in providing the annotations. Therefore, we argue that TaxVAMB provide high-quality taxonomic annotations for the bins created from data from well-studied environments without the need to use additional MAG taxonomic classification tools.

### TaxVAMB uncovers both bacterial and fungal MAGs in wheat phyllosphere dataset

Finally, we investigated the results of applying TaxVAMB to the short-read wheat phyllosphere dataset. The dataset consisted of 211 samples from the surface of wheat flag leaves at nine time points during the end of growth season of 2022 from a single field in Denmark (see Methods). From the dataset we reconstructed 614 high quality and 647 medium quality bacterial MAGs across five phyla (Actinomycetota, Bacillota, Bacteroidota, Deinococcota, Pseudomonadota) (Figure 6a, Supplementary Figure S9). We found that the binned MQ and HQ MAGs explained 13.4%-98.4% of the total reads across the samples with a mean of 49.2% of the reads (Figure 6c). When investigating the prevalence of the species, we found that the five most prevalent species measured in the number of MAGs assigned were present in 30-60% of all the samples (*Pseudomonas poae, Frigoribacterium sp001421165, Pseudomonas graminis, Pantoea agglomerans, Erwinia aphidicola*) and had previously been described in literature as part of the wheat phyllosphere ^58–63^ (Figure 6b, Supplementary Table 2, Supplementary Table 3). In addition to these, TaxVAMB reconstructed a novel species of genus Sphingomonas that was present in 12% of samples with an abundance of >1% of the mapped reads. Additionally, we discovered that the *Pantoea agglomerans* species was more prevalent as the plants senesced (Mann-Whitney U test, p-value = 2e-18) (Supplementary Figure S9). We also tested the ability of TaxVamb to recover fungal bins by investigating the bins that were annotated by TaxVAMB as Eukaryotes.

**Fig. 6.**
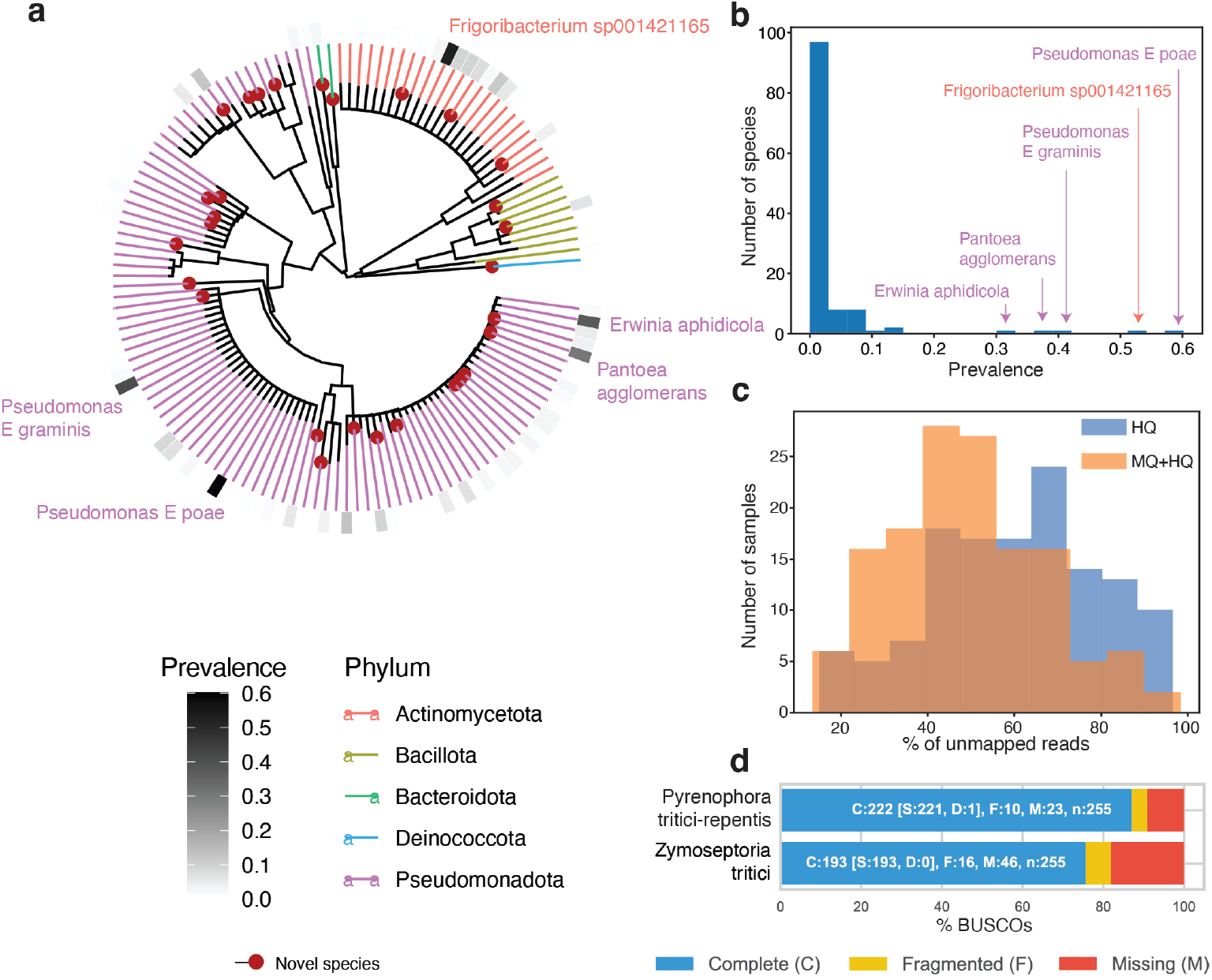
Wheat phyllosphere MAGs. **a**. Phylogenetic tree of high quality bacterial MAGs indicating the most prevalent species in terms of high quality MAGs per sample. **b**. Distribution of prevalences for all species. The top 5 most prevalent species are annotated with labels. **c**. Distributions of shares of unmapped reads for each sample with MQ or HQ MAGs. Blue color (HQ) is the share of unmapped reads when only mapping to HQ MAGs (completeness > 90%, contamination < 5%). Orange color (HQ+MQ) is the share of unmapped reads when mapping to HQ and MQ MAGs (completeness *≥* 50%, contamination < 10%). **d**. BUSCO results for two fungal MAGs, annotated by TaxVAMB as *Zymoseptoria tritici* and *Pyrenophora tritici-repentis*.

Two of such bins were larger than 27Mb, with 99.9% and 20% of their contigs annotated with fungal species after the Taxometer step. Thus, we recovered *Zymoseptoria tritici* and *Pyrenophora tritici-repentis* MAGs with corresponding BUSCO ^64^ completeness scores of 75% and 87% (Figure 6d). Both of these fungal species are known wheat pathogens ^65,66^. We conclude that TaxVAMB recovered a large variety of novel MAGs of medium and high quality, providing insights into the bacterial and fungal composition of the wheat phyllosphere.

## Discussion

In summary, we present TaxVAMB, a method for combining intrinsic features (TNFs and abundances) with annotation features (taxonomic labels). We utilize the full hierarchical structure of taxonomic labels with a deep hierarchical loss, allowing us to train the model even on contig annotations from higher taxonomic ranks. We justified the network architecture choice by empirically evaluating a competing stacked VAE model. The labels were also used to assign the preliminary taxonomic annotations to MAGs down to the species level, matching GTDBtk tool in performance for the high quality bins when tested on CAMI2. In the number of high quality bins, Tax-VAMB exceeds or matches the performance of other binners for the genomes that are almost completely present in the input, and shows unmatched performance in binning incomplete genomes.

We identified two conditions where TaxVAMB exhibits the biggest gains compared to previous state-of-the-art: a) sufficiently high quality taxonomic labels which occurred for the datasets from more well studied environments such as human gut; b) a relatively low number of samples where the signal from the co-abundance vector was less strong (< 100 samples judging from the Figure 5). For example, we expect that one use case of TaxVAMB is application to studies of the human microbiome where few samples are available. Similarly, TaxVAMB also shows unmatched performance for binning incomplete genomes, which, just like datasets with a small number of samples, produce low quality abundance vectors. We notice that using TaxVAMB in poorly studied and much more complex environments such as anaerobic digester sludge does not degrade the performance compared to the unsupervised binning provided by VAMB. We also demonstrated that TaxVAMB does not rely on single-copy genes for reaching optimal performance, which allows it to better bin incomplete genomes and non-bacterial entities.

Predicting annotations for the full dataset based on pre-annotated contigs might increase the bias toward well-studied taxa, since these taxa are more likely to be returned by a taxonomic classifier such as MMseqs2. The qualities of predictions depend on the share of annotated contigs and taxonomic diversity of the samples. We address the possible bias in two ways. First, the taxonomy predictions come as probability vectors for each taxonomic rank, allowing to set a confidence threshold for predictions. A contig with an ambiguous prediction will remain unannotated on this taxonomic rank and below. Second, unsupervised learning still allows correct binning of contigs without any good match in the database, but which share intrinsic features such as TNFs and abundances with each other.

The databases of reference genomes are constantly updated (GTDB increased by around 30% in terms of bacterial species clusters from v207 to v214 ^67^, NCBI estimates annual growth in terms of the number of genomes at 15% ^68^). The bias introduced by taxonomic annotation of a subset of contigs in a dataset will continue to reduce as the number of diverse genomes in databases grows.

After reclustering using single-copy genes, unsupervised (VAE) and self-supervised (contrastive learning) models demonstrate similar performance on short- and long-read datasets. While we flag optimizing for single-copy genes metrics during training as a potential source of error when also used as an evaluation metric, we also interpret the large gains from single-copy gene-guided reclustering as the indicator that future research in the field of metagenomic binning would find the biggest gains in integrating new data modalities with intrinsic features, and not solely in applying new algorithms to TNFs and abundances.

Multi-omics data integration is a powerful technique for understanding complex biological systems ^69–71^. Semi-supervised multimodal VAEs can be easily adapted to learn from weakly labeled heterogeneous multi-omics datasets in other fields beyond metagenomics binning. The same applies to the hierarchical loss, since many biological data types follow a hierarchical structure.

Improving over previous attempts to incorporate taxonomic labels to the metagenome binning process, we conclude that it greatly improves binning when the full hierarchical structure is in use, and is a great additional input when the abundance vector is not informative enough. As the quality of reference databases will improve over time, the impact of using TaxVAMB will be potentially even more powerful in the future.

## Methods

### Bi-modal variational autoencoder

VAE is a generative model performing variational inference over the latent variable *z*. The model is formally defined as *p*(*x, z*) = *p*(*z*)*p*(*x*|*z*). The intractable posterior *q*(*z*|*x*) and the conditional distribution *p*(*x*|*z*) are approximated by neural networks using the ELBO-loss function:

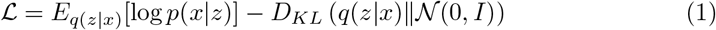

The bi-modal VAE extends the basic VAE by allowing training and inference on the dataset where: a) the input consists of two modalities and b) a modality can be missing for one or more samples. We define the modality here as the part of the data that is missing for part of the dataset. Thus notice that while the VAMB model is trained on both TNFs and abundances, we do not define it as bi-modal for the purpose of this summary, since both TNFs and abundances are present for all samples and can be trivially converted into one modality by concatenating the corresponding input vectors.

While the VAE approximates the posterior *q*(*z*|*x*) with a neural network encoder that takes *x* as an input, bi-modal VAE extends this approach by modelling *q*(*z*|*x*_1_, *x*_2_), *q*_1_(*z*|*x*_1_) and *q*_2_(*z*|*x*_2_), which replace the single *q*(*z*|*x*). There are two decoders approximating distributions *p*(*x*_1_|*z*) and *p*(*x*_2_|*z*). Multimodal VAEs differ in 1) the way they approximate *q*(*z*|*x*_1_, *x*_2_), *q*_1_(*z*|*x*_1_) and *q*_2_(*z*|*x*_2_) by neural networks and/or 2) the structure of the loss function.

TaxVAMB implements the VAEVAE ^50^ model from bi-modal VAE family which models *q*(*z*|*x*_1_, *x*_2_), *q*_1_(*z*|*x*_1_) and *q*_2_(*z*|*x*_2_) by corresponding neural networks. The following ELBO-like loss L is minimised:

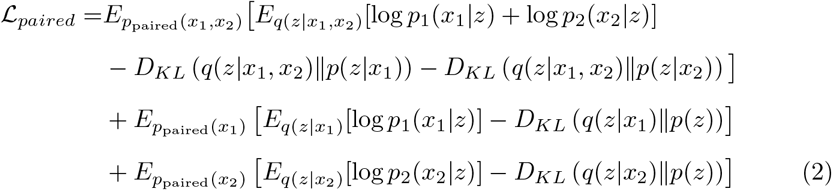

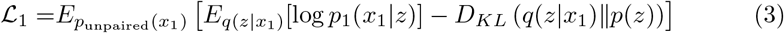

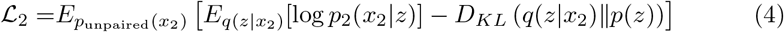

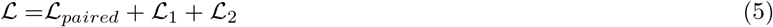

with *D*_*KL*_ (*p*(*x*)∥*q*(*x*)) being the Kullback–Leibler divergence between two probability distributions *p*(*x*) and *q*(*x*), defined as:

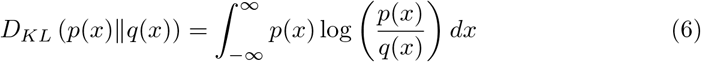

The training procedure includes constructing the dataset with paired and unpaired samples. Let *C* be a list of all contigs. Three copies of *C*, denoted as *C*_*paired*_, *C*_1_, and *C*_2_, are independently shuffled. The paired samples are ordered tuples (*x*_1_, *x*_2_) where *x*_1_ is a concatenation of TNF vector and abundance vector (the input of VAMB) and *x*_2_ is a taxonomy label vector described in Subsection 6, and *x*_1_ and *x*_2_ correspond to the same contig from the set *C*_*paired*_. An unpaired TNFs and abundances vector 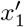 corresponds to a contig from the list *C*_1_. An unpaired taxonomy label corresponds to a contig from the list *C*_2_. The forward pass follows the steps from the Algorithm 1.

#### Algorithm 1 Loss computation (forward pass)

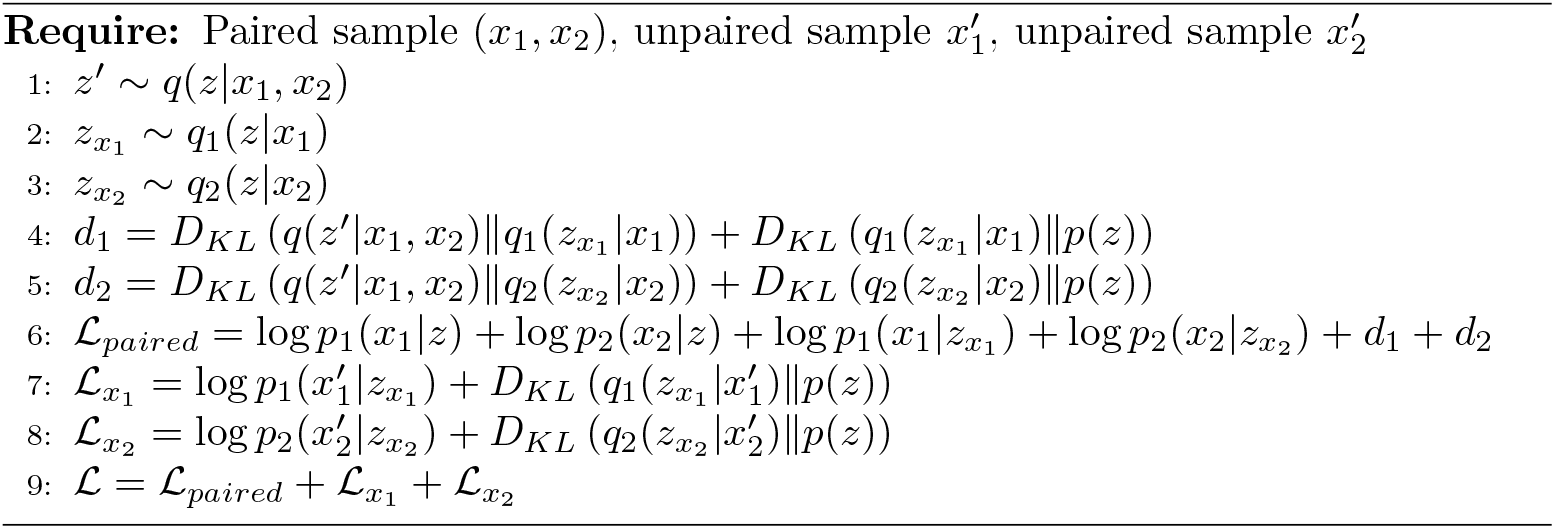

### Data preprocessing

The workflow of preprocessing the data is the same as in Taxometer (v.b5fd0ea) ^42^ and VAMB ^11^. The synthetic short paired-end reads from each sample were aligned using bwa-mem (v.0.7.15) ^72^. BAM files were sorted using samtools (v.1.14) ^73^. Contigs ≤2,000 base pairs (bp) were removed for each dataset. the long-read datasets were both sequenced using Pacific Biosciences HiFi technology. We assembled each sample using metaMDBG (v.b55df39) ^74^, mapped reads using minimap2 (v.2.24) ^75^ with the ‘-ax map-hifi’ setting, and then continued with the same workflow as with the short reads.

### Abundances and TNFs

The workflow of computing abundances and TNFs is the same as in Taxometer (v.b5fd0ea) ^42^ and VAMB ^11^. Computation of abundances and TNFs was done using the VAMB metagenome binning tool ^11^. To determine TNFs, tetramer frequencies of non-ambiguous bases were calculated for each contig, projected into a 103-dimensional orthonormal space and normalized by z-scaling each tetranucleotide across the contigs. To determine the abundances of each sample, we used pycoverm 0.6.0 ^76^. The abundances were first normalized within sample by total number of mapped reads, then across samples to sum to 1. To determine absolute abundance, the sum of abundances for a contig was taken before the normalization across samples. The dimensionality of the feature table was then *N*_*c*_ × (103 + *N*_*s*_ + 1) where *N*_*c*_ was the number of contigs, *N*_*s*_ was the number of samples.

### Network architecture and hyperparameters

The encoder architectures for the concatenated vector of abundances and TNFs is the same as in Taxometer (v.b5fd0ea) ^42^ and VAMB ^11^. The input vector of dimensionality *N*_*c*_ × (103 + *N*_*s*_ + 1) was passed through 4 fully connected layers ((103 + *N*_*s*_ + 1) × 512, 512 × 512, 512 × 512, 512 × 512) with leaky ReLU activation function (negative slope 0.01), each using batch normalization (epsilon 1*e* − 05, momentum 0.1) and dropout (*P* = 0.2).

The encoder network for the taxonomy labels had the input dimensions of *N*_*l*_ where *N*_*l*_ was the number of leaves in the taxonomic tree. The input vector was passed through 4 fully connected layers (*N*_*l*_ × 512, 512 × 512, 512 × 512, 512 × 512) with leaky ReLU activation function (negative slope 0.01), each using batch normalization (epsilon 1*e* − 05, momentum 0.1) and dropout (*P* = 0.2).

The encoder network for the concatenation of the two modalities had the input dimensions of (103 + *N*_*s*_ +1) + *N*_*l*_ where *N*_*s*_ was the number of samples and *N*_*l*_ was the number of leaves in the taxonomic tree. The input vector was passed through through 4 fully connected layers ((103 + *N*_*s*_ + 1) × 512, 512 × 512, 512 × 512, 512 × 512) with leaky ReLU activation function (negative slope 0.01), each using batch normalization (epsilon 1*e* − 05, momentum 0.1) and dropout (*P* = 0.2).

The bi-modal VAE has two decoder networks, one for each modality. Both of them follow the same architectures as the corresponding encoders, with the input vector having the dimensionality of the latent space, and the output having the dimensionality of the corresponding modality.

For short-read datasets, the network was trained for 300 epochs with batch size 256, latent space dimensionality 32. For long-read datasets, the network was trained for 1000 epochs with batch size 1024, latent space dimensionality 64. All models were using the Adam optimizer with learning rates set via D-Adaptation ^77^. The model was implemented using PyTorch (v.1.13.1) ^78^, and CUDA (v.11.7.99) was used when running on a V100 GPU.

### Hierarchical loss

The hierarchical loss is the same as in Taxometer (v.b5fd0ea) ^42^. A phylogenetic tree was constructed for each dataset from the taxonomy classifier annotations for the set of contigs. Thus, the resulting taxonomy tree *T* was a subgraph of a full taxonomy and the space of possible predictions was restricted to the taxonomic identities that appeared in the search results. For the above experiments, we used a flat softmax loss. Let *N*_*l*_ be the number of leaves in the tree *T*. The likelihoods of leaf nodes of the taxonomy tree were obtained from the softmax over the network output layer with dimensionality 1 ×*N*_*l*_. The likelihood of an internal node was then a sum of likelihoods of its children and computed recursively bottom-up. The model output was a vector of likelihoods for each possible label. For the backpropagation, the negative log-likelihood loss was computed for all the ancestors of the true node and the true node itself. Predictions were made for all taxonomic levels and for each level, the node descendant with the highest likelihood was selected. If no node descendant had likelihood > 0.5, the predictions from this level and the levels below were not included in the output.

### Taxonomic classifiers

We obtained the taxonomic annotations for contigs of all seven short-read and two long-read datasets from MMseqs2 (v.7e2840) ^33^, Metabuli (v.1.0.1) ^79^, Centrifuge (v.1.0.4) ^80^ and Kraken2 (v.2.1.3) ^81^. For MMseqs2, we used the mmseqs taxonomy command. For Metabuli, we used the metabuli classify command with *–seq-mode 1* flag. For Centrifuge, we used the centrifuge command with *-k 1* flag. For Kraken2, we used the kraken command with *–minimum-hit-groups 3* flag. MMseqs2 and Metabuli were configured to use GTDB v207 as the reference database. Centrifuge, Kraken2 and MetaMaps were configured to use NCBI identifiers. All the taxonomic annotations were first refined with Taxometer ^42^ (v.b5fd0ea) with the default parameters (epochs 100, batch size 1024).

### Benchmarked binners

Metabat (v.2.12.1) *metabat* command with the default parameters is used. Metadecoder (v.1.0.19) *coverage, seed* and *cluster* commands are used as described in https://github.com/liu-congcong/MetaDecoder. Comebin (v.1.0.3) *run comebin*.*sh* command with the default parameters is used. SemiBin2 (v.2.1.0) *multi easy bin* command is used with the flags *–engine gpu, –separator C, -t 20, –write-pre-reclustering-bins* and *–self-supervised*. VAMB, AVAMB and TaxVAMB were run as a part of the VAMB codebase (commit 5f2cd7), with the corresponding commands *”vamb bin default”, “vamb bin avamb”* and *”vamb bin taxvamb”*.

### Reclustering using SCGs

Short-read and long-read reclustering algorithms that use single-copy genes are the same as introduced in SemiBin2 ^82^. The code was adapted from the SemiBin2 code-base (https://github.com/BigDataBiology/SemiBin/blob/main/SemiBin/long_read_cluster.py and https://github.com/BigDataBiology/SemiBin/blob/main/SemiBin/cluster.py) for the TaxVAMB codebase (https://github.com/RasmussenLab/vamb/blob/master/vamb/reclustering.py). TaxVAMB uses the same 107 single-copy marker genes as used in the SemiBin2 tool to estimate the completeness, contamination, and F1-score of every bin. Completeness for each bin is calculated as 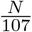, contamination as 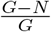 and F1-score as 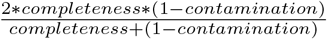. *N* is the number of different single-copy genes in a bin, *G* is the total number of sequences matching any single-copy gene.

For the short-read datasets, k-means based reclustering of TaxVAMB/VAMB clusters is performed. Bins where more than one marker gene of the same kind is present are reclustered with the weighted k-means method using the contigs containing the repeated marker gene as the initial centroids. It results in bins with reduced contamination.

For the long-read datasets, the DBSCAN algorithm from a Python library *scipy* (v1.10.0) was used to perform the clustering from scratch (the previous clusters, made by TaxVAMB/VAMB, were not used). Same as in SemiBin2, DBSCAN was run with *ϵ* value equals to 0.01, 0.05, 0.1, 0.15, 0.2, 0.25, 0.3, 0.35, 0.4, 0.45, 0.5, and 0.55. From all the resulting bins, the best one by the F1-score was recursively selected, and its contigs were removed from all the remaining bins, after which the selection of the best bin repeats. It repeats until no more bins fulfill the criteria for minimal quality (completeness more than 90%, contamination less than 5%). One change that was made in the TaxVAMB long-read reclustering was that it performed the described procedure per a set of contigs assigned to the same genus by the Taxometer refinements of the provided taxonomic annotations.

### CAMI2 benchmarks

For short-read benchmarking, we used five CAMI2 datasets: Airways (10 samples), Oral (10 samples), Skin (10 samples), Gastrointestinal (10 samples), Urogenital (9 samples), assemblies for which were sample-specific. We benchmarked the following binners on the synthetic CAMI2 toy human microbiome dataset: Metabat ^10^, MetaDecoder ^83^, COMEBin ^22^, SemiBin2 ^82^, AVAMB ^28^, and VAMB ^11^. We used taxonomic labels from four taxonomic classifiers as an input to TaxVAMB: MMSeqs2 ^33^, Metabuli ^79^, Kraken2 ^81^ and Centrifuge ^80^. AVAMB, VAMB and TaxVAMB bins were postprocessed with reclustering using single-copy genes. We used the number of high quality bins and assemblies estimated via BinBencher (v0.3.0) ^54^ as a metric. For the MAG taxonomic annotation experiment we used CheckM2 (v1.0.2) ^40^.

We benchmarked using BinBencher (v.0.3.0) ^54^ against a reference computed from the CAMI2 ground truth. The metrics used were number of near-complete (defined as recall ≥ 0.9, precision ≥ 0.95) assemblies or genomes. As defined in the BinBencher paper, for genomes, the recall was counted relative to the full length of the genome from which the reads were simulated from, whereas when counting assemblies, the recall was relative to the assembled part of those genomes, i.e. the part of the genomes covered by a contig which was used as input to the binner. The number of near-complete genomes reflect the MAG quality relative to the underlying biological organism and thus depends more on limitations of the dataset, whereas the assembly metric may better reflect the methodological gains from using a different algorithm.

### Long-read benchmarks

For long-read benchmarking we used a human gut microbiome dataset with 4 samples and a dataset from anaerobic digester sludge with 3 samples ^84^, both sequenced using Pacific Biosciences HiFi technology. We assembled each sample using metaMDBG (v.b55df39) ^74^, mapped reads using minimap2 (v.2.24) ^75^ with the ‘-ax map-hifi’ setting, and from there proceeded as with the short reads. For evaluating the quality (completeness and contamination) of the resulting MAGs we used CheckM2 (v.1.0.2) ^40^. The Metadecoder binner failed on the human gut dataset experiment with an internal code error and its results are thus not displayed on the figure.

### Multisample scaling

For the experiment that evaluates the number of bins given a different number of samples, we used a short-read human gut dataset with 1,000 samples from Almeida et al. ^57^, as well as our own wheat phyllosphere dataset with 211 samples. For each dataset, we split all the samples into three sets of chunks: 1) chunks of 100 samples; 2) chunks of 10 samples; 3) chunks of 1 sample. Each chunk was treated as an independent dataset We then summed the resulting number of near-complete bins within each set of chunks.

### Wheat phyllosphere dataset: sample collection and processing

Twenty-four field plots of *Triticum aestivum* were sampled by collecting composite samples of 30 flag leaves nine times between June 7th and July 14th 2022, at a field trial in Ringsted, Denmark. The experimental design included three wheat cultivars, four replicates, and two treatments, which were unsprayed and sprayed with a fungicide. The samples were washed in 100 ml wash solution (0.9% NaCl + 0.05% Tween80), vigorously shaken for 2 minutes, sonicated for 2 minutes and then vigorously shaken again for 2 minutes, filtered (10 µm), centrifuged (4000 × g, 15 min) and the pellet resuspended in 1 ml 1x PBS and stored at -20°C until DNA extraction using the FastDNA™ SPIN Kit (MP Biomedicals, CA, USA) for Soil according to instructions eluding in 100 µl DES. DNA libraries were build using the Illumina Nextera XT kit (Illumina, CA, USA), but samples with <0,1 ng/µl DNA were built with a 1-fold diluted ATM, 20 PCR cycles and a higher ratio of AMPure XP beads (Beckman Coulter, IN, USA) ^85^. Libraries were sequenced using Illumina paired-end (2 x 150bp) technology (NovaSeq 6000 S4 v1.5).

### Wheat phyllosphere dataset: data analysis

Raw sequencing reads were filtered using fastp (v.023.2) ^86^ with the following option: `‘--trim_tail 1 --cut_tail --trim_poly_g --dedup --length_required 80’`. Quality control of the filtered reads was assessed using MultiQC (v.1.12) ^87^. To remove reads originating from wheat or potential human contamination, the reads were mapped to the reference sequences GCF 018294505.1, MG958554.1 and GCF 000001405.40 (GRCh38.p14). Mapping was performed using Bowtie ^88^(v.2.5.3). Paired reads where both mates were unmapped were extracted using Samtools (v.1.18) ^73^ with the `‘fastq -f 13’` option.

Metagenomic assemblies were generated for each sample using SPAdes (v.3.15.4) ^89^ with the `‘--meta -k 21,33,55,77,99’` option. Assembly statistics were computed using QUAST (v.5.2.0) ^90^.

MAGs are assigned the taxonomy using GTDBtk (v.2.4.0) configured with the GTDB database v220. Taxonomic trees are built using *ggtree* ^91^(v.3.19), *tidytree* (v.0.4.6), *treeio* R (v.4.4.1) libraries. Mann-Whitney U test is performed on *Pantoea agglomerans* abundances by splitting the samples into two groups: 143 samples from the earlier days (2022-06-07, 2022-06-10, 2022-06-14, 2022-06-17, 2022-06-21) and 103 samples from the later days (2022-06-28, 2022-07-04, 2022-07-07, 2022-07-14) using *scipy* (v1.10.0).

## Acknowledgements

S.K., M.N., S.R., N.S.O., L.R., L.M.F.J., P.E.D., A.G., K.N.N., S.C. and L.H. were supported by the Novo Nordisk Foundation (grant NNF19SA0059348). P.P., J.N.N., S.K., K.N.N. and S.R. were supported by the Novo Nordisk Foundation (grant NNF23SA0084103). S.K., P.P., J.N.N., K.N.N. and S.R. were supported by the Novo Nordisk Foundation (grant NNF14CC0001). P.P., J.N.N. and S.R were supported by the Novo Nordisk Foundation (grant NNF20OC0062223). S.R. was supported by the Novo Nordisk Foundation (grant NNF21SA0072102). We thank Chayan Roy, Sif Christine Lykke Hougaard and Xianfu Liu for contributing to the wheat phyllosphere data collection.

## Author Contributions

S.K., M.N., J.N.N. and S.R. designed the experiments. P.P., J.N.N. and K.N.N. preprocessed the datasets. S.K. wrote the software and performed the analysis. M.N., P.P., J.N.N. and S.R. provided guidance and input for the analysis. S.C. and J.C.W. selected the trial fields and developed sampling protocols for the wheat phyllosphere dataset. N.S.O., L.R., L.M.F.J., P.E.D., A.G., K.N.N., S.C. and L.H. performed sample collection, sample processing, DNA extractions and library building for the wheat phyllosphere dataset. S.K. wrote the manuscript with contributions from all coauthors. All authors read and approved the final version of the manuscript.

## Data availability

The CAMI2 datasets were downloaded from https://data.camichallenge.org/participate from “2nd CAMI Toy Human Microbiome Project Dataset” (5 human microbiome datasets), “2nd CAMI Challenge Marine Dataset” (Marine), “2nd CAMI Challenge Rhizosphere challenge” (Rhizosphere). The long-read human gut dataset is available at https://downloads.pacbcloud.com/public/dataset/Sequel-IIe-202104/metagenomics/. The long-read sludge dataset is available at the ENA as part of the study PRJEB39861. The 1000 samples short-read human gut dataset was first published by Almeida et al. The de novo assemblies of the Almeida dataset were obtained through personal communication with A. Almeida and R. D. Finn, and the reads downloaded from ENA ERP108418 as specified in their publication. The phyllosphere short-read dataset is available at the ENA using the accession ERP165292. The near complete and medium quality MAGs from the phyllosphere are available for download at Zenodo ^92^ by the link https://zenodo.org/records/13959411. The MAGs will be uploaded to ENA upon acceptance as they might change during revisions.

## Code availability

All code can be found on GitHub at https://github.com/RasmussenLab/vamb and is freely available under the permissive MIT license. The code for making the figures is in the separate repository on Github https://github.com/sgalkina/TaxVAMB_paper_figures.

