## Supplementary Material for "Binning meets taxonomy: TaxVAMB improves metagenome binning using bi-modal variational autoencoder"

### Supplementary materials for the paper "Binning meets taxonomy: TaxVAMB improves metagenome binning using bi-modal variational autoencoder"

#### List of Figures

S1 Stacked VAE, Siamese networks and bi-modal VAE architectures . . . S3
S2 Shotgun metagenomics computational workflows without and with TaxVAMB . . . S4
S3 Metagenome binners performance measured in high quality assemblies S5
S4 Metagenome binners performance measured in high quality genomes . S6
S5 Metabuli vs TaxVAMB . . . S7
S6 Bi-modal VAE vs stacked VAE performance . . . S8
S7 Taxonomic annotations for MAGs . . . S9
S8 Majority vote taxonomic annotation of high quality bins for TaxVAMB and VAMB . . . S10

S9 Abundance matrix for all species and samples, wheat phyllosphere. . . S11

### List of Tables

1 Metagenome binners runtimes . . . . . . . . . . . . . . . . . . . . . . . S12

2 Wheat phyllosphere species prevalence . . . . . . . . . . . . . . . . . . . S13

3 Wheat phyllosphere genus prevalence . . . . . . . . . . . . . . . . . . . . S14

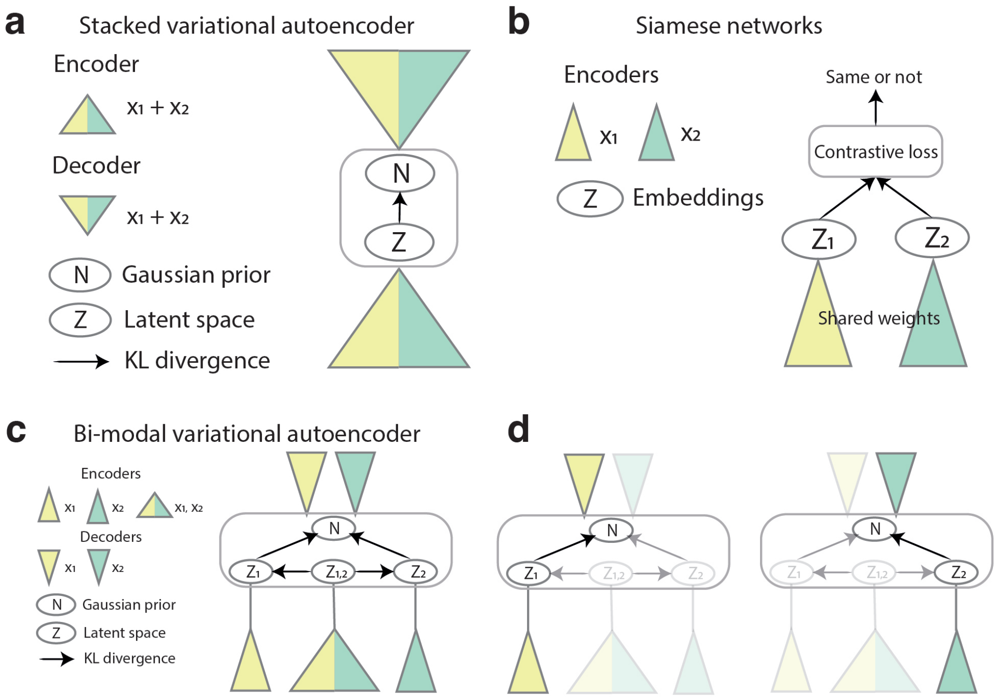


**Supplementary Figure S1 Stacked VAE, Siamese networks and bi-modal VAE architectures.** The input in on the bottom, the output is on the top. **a.** Stacked variational autoencoder, with the concatenated vector of the two modalities as an input and output to the basic VAE architecture. **b.** Siamese networks, with two modalities as separate inputs to the two encoders, and the shared weights between the two encoders. **c.** Bi-modal variational autoencoder architecture, with the two encoders for each modality and one joint encoder for the concatenated vector of the two modalities. **d.** The input in the training and the infrerence time when only one of the modalities is present.

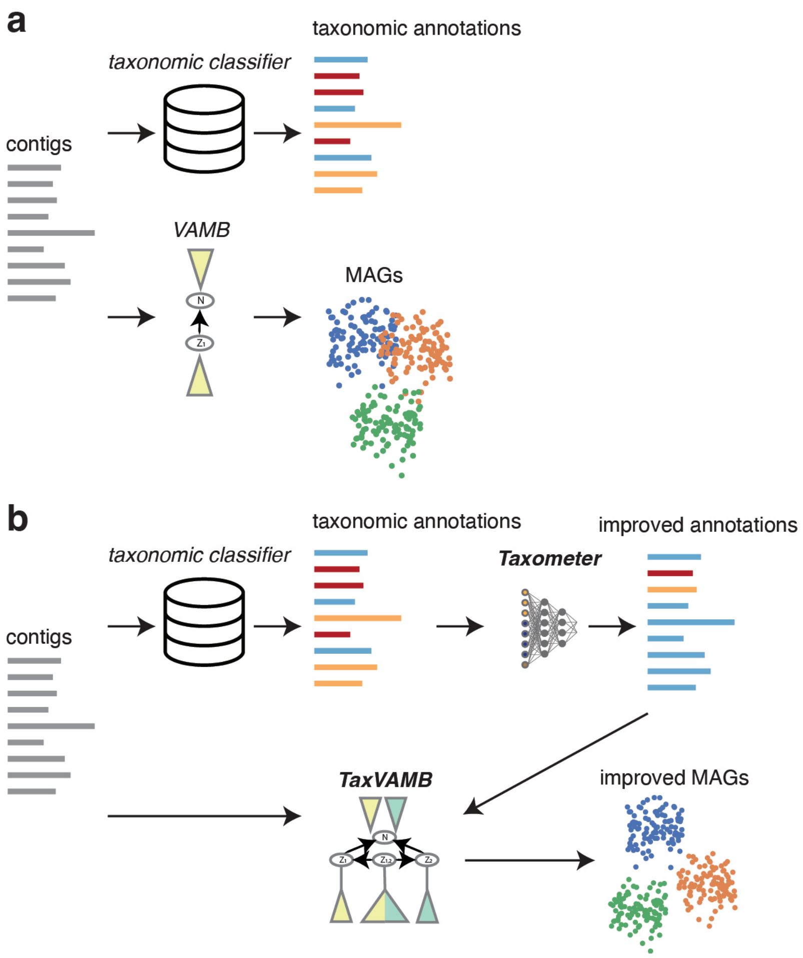


**Supplementary Figure S2 Metagenomic analysis workflows without and with TaxVAMB. a.** Without TaxVAMB, the taxonomic classification of contigs and the metagenome binning are performed as two independent workflows. **b.** With TaxVAMB, taxonomic annotations from a taxonomic classifier are first refined using Taxometer, and then used as the second modality in the bi-modal VAE, resulting in higher quality MAGs.

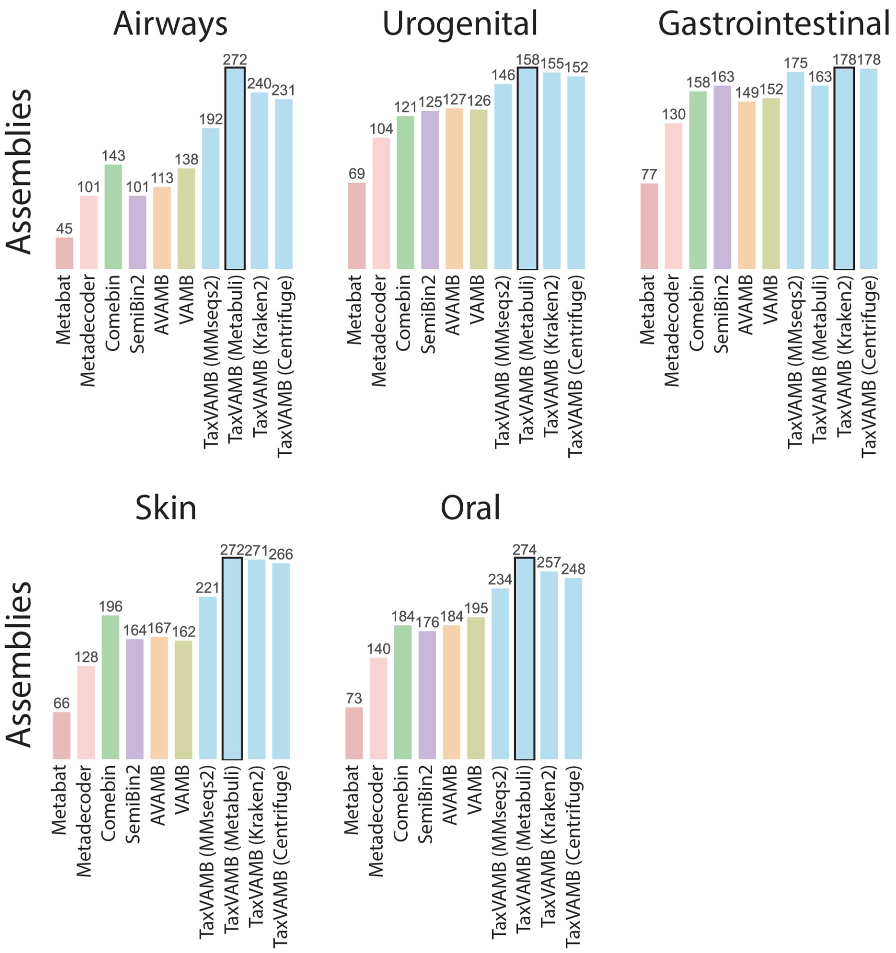


**Supplementary Figure S3 Metagenome binners performance measured in high quality assemblies.** Metagenome binners benchmarks on CAMI2 human microbiome datasets, with TaxVAMB using four different taxonomic classifiers. The metrics is the number of high quality assemblies. The assembly metrics measures recall in respect to the part of the genome provided to the binner. SemiBin2, VAMB and TaxVAMB results are after applying the k-means based reclustering step.

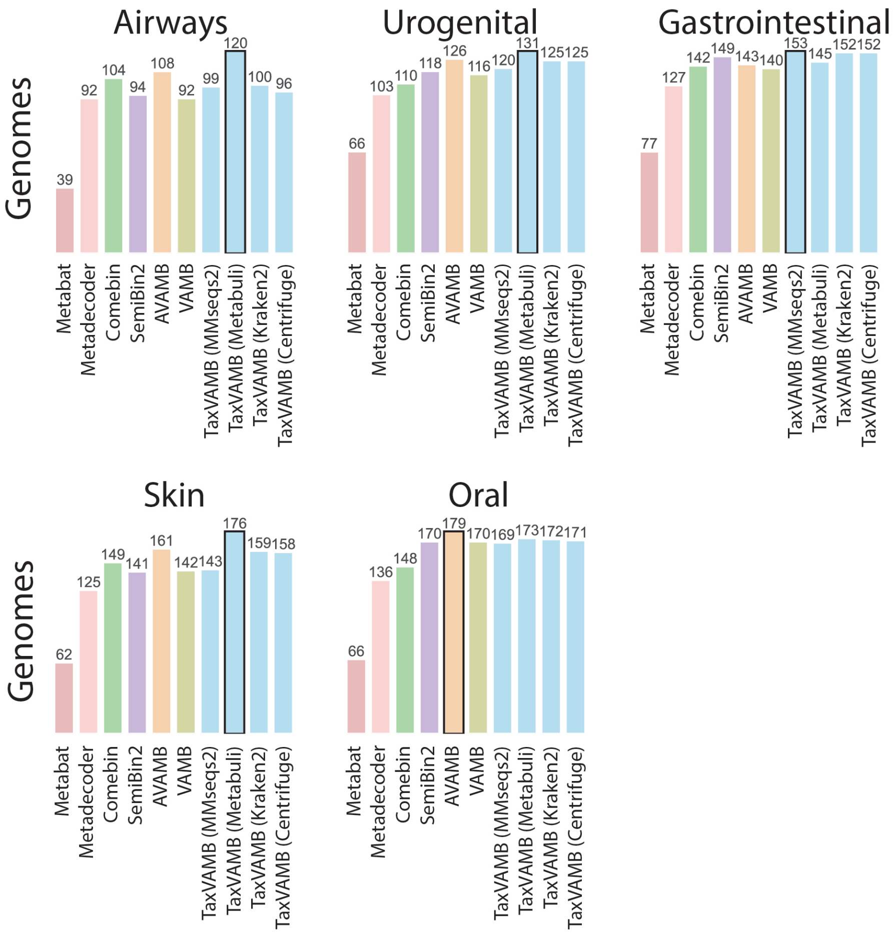


**Supplementary Figure S4 Metagenome binners performance measured in high quality genomes.** Metagenome binners benchmarks on CAMI2 human microbiome datasets, with TaxVAMB using four different taxonomic classifiers. The metrics is the number of high quality full genomes. The full genome metrics measures recall in respect to the full genome. SemiBin2, VAMB and TaxVAMB results are after applying the k-means based reclustering step.

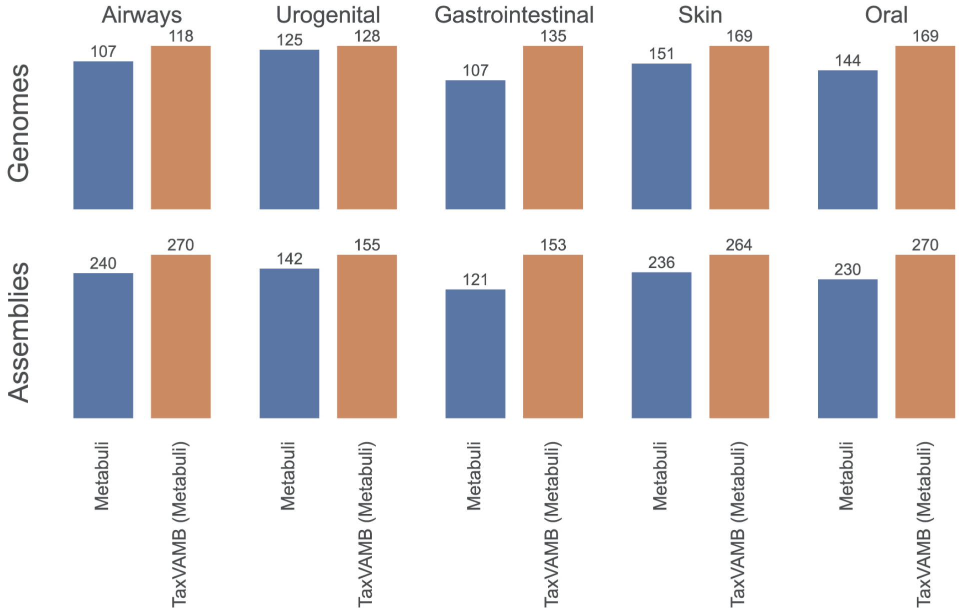


**Supplementary Figure S5 Metabuli vs TaxVAMB.** Since Metabuli classifier outputs labels down to the subspecies level, the contigs that were assigned the same subspecies label can be used as a MAG. Here, Matebuli is compared to TaxVAMB that uses Metabuli labels on the CAMI2 toy datasets using the high quality genomes and the high quality assemblies metrics.

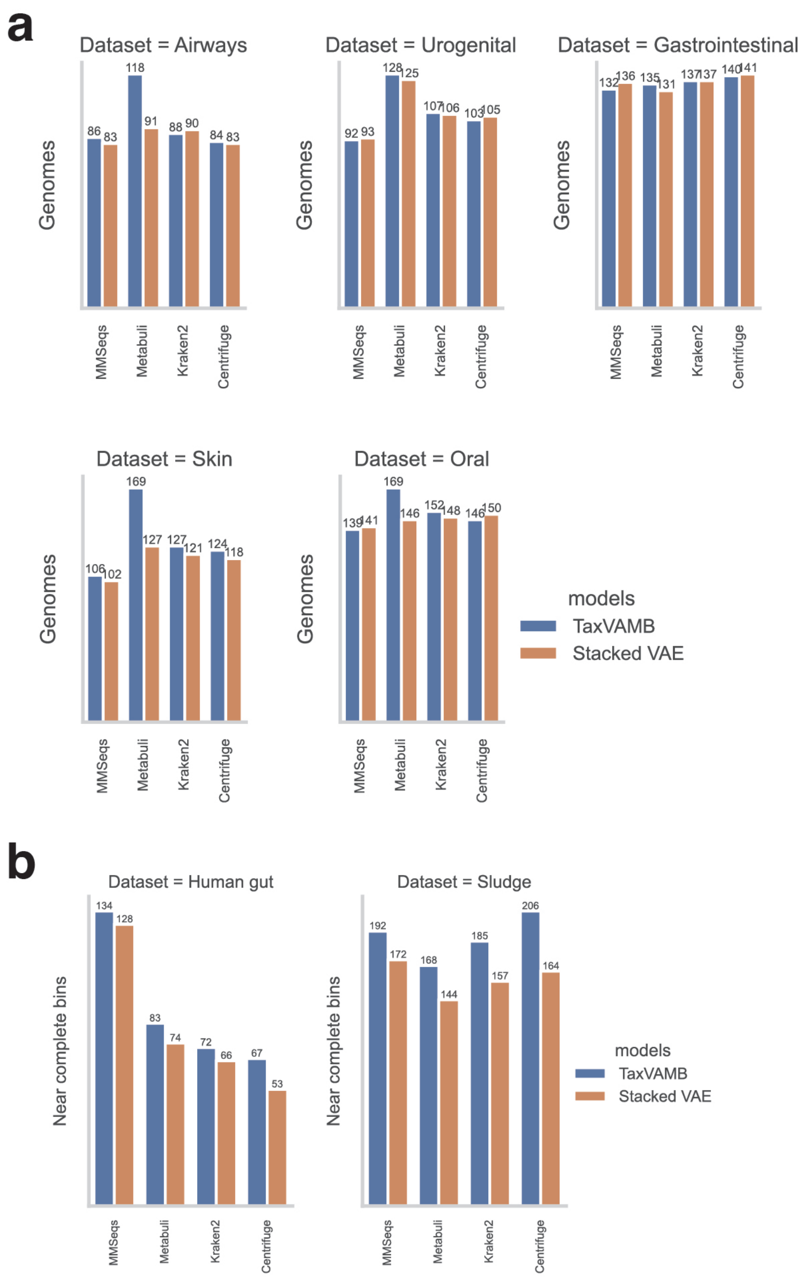


**Supplementary Figure S6 Bi-modal VAE vs stacked VAE performance.** Comparison of the number of high quality genomes as estimated by BinBencher (CAMI2 datasets) and CheckM2 (long-read Human Gut and Sludge datasets) for TaxVAMB presented architecture (bi-modal VAE) and the modification of TaxVAMB to use the stacked VAE for processing the two modalities.

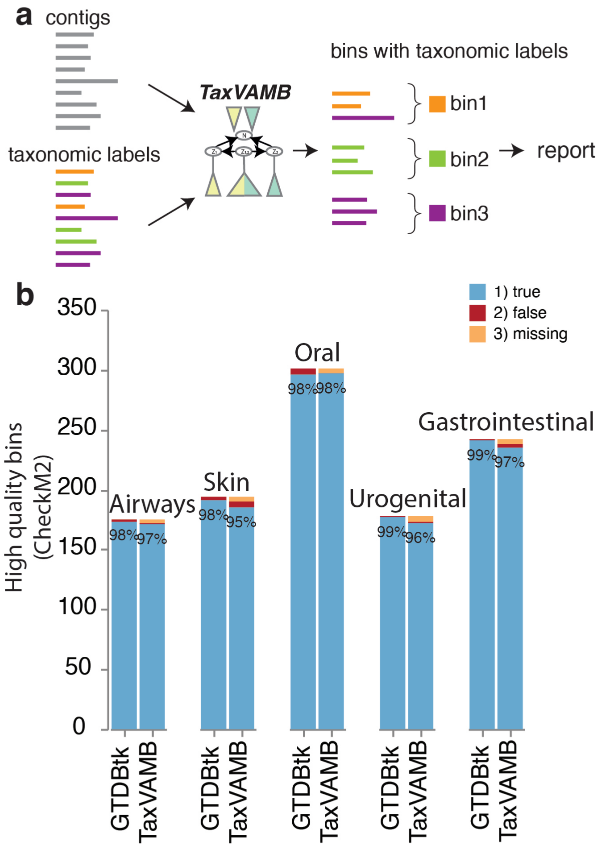


**Supplementary Figure S7 MAGs with taxonomic annotations. a.** The TaxVAMB output, resulting in preliminary taxonomic classifications for all MAGs. **b.** The evaluation of the quality of the MAG taxonomic assignments for CAMI2 datasets, based on the provided ground truth labels, compared to GTDB-tk classification results for high quality bins, as evaluated by CheckM2. Kraken2 taxonomic labels were used.

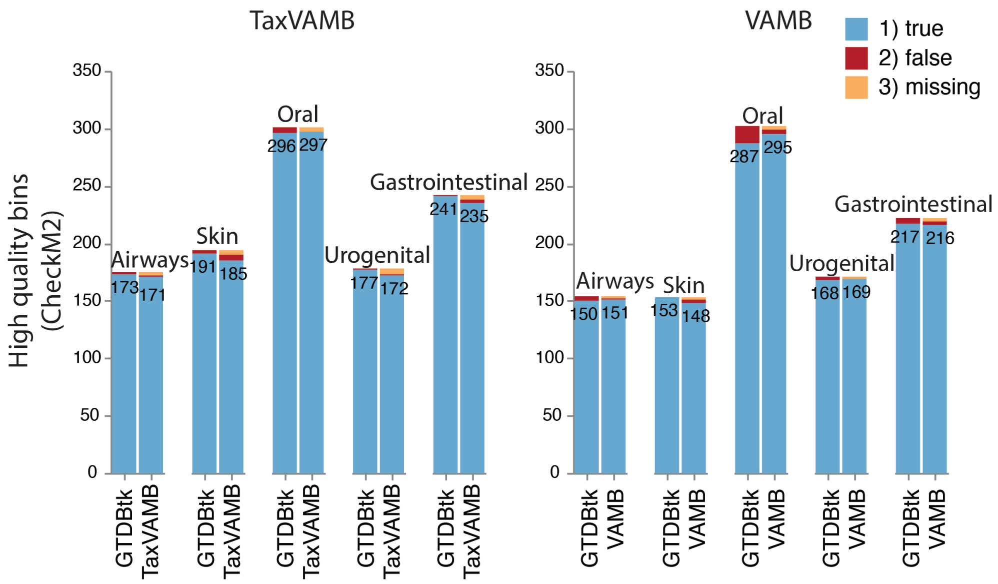


**Supplementary Figure S8 Majority vote taxonomic annotation of high quality MAGs for TaxVAMB and VAMB.** High quality MAGs of TaxVAMB and VAMB (evaluated via CheckM2) are taxonomically classified by GTDB-tk software and by majority vote of contigs taxonomic assignment from Kraken2 for each bin for CAMI2 toy datasets.

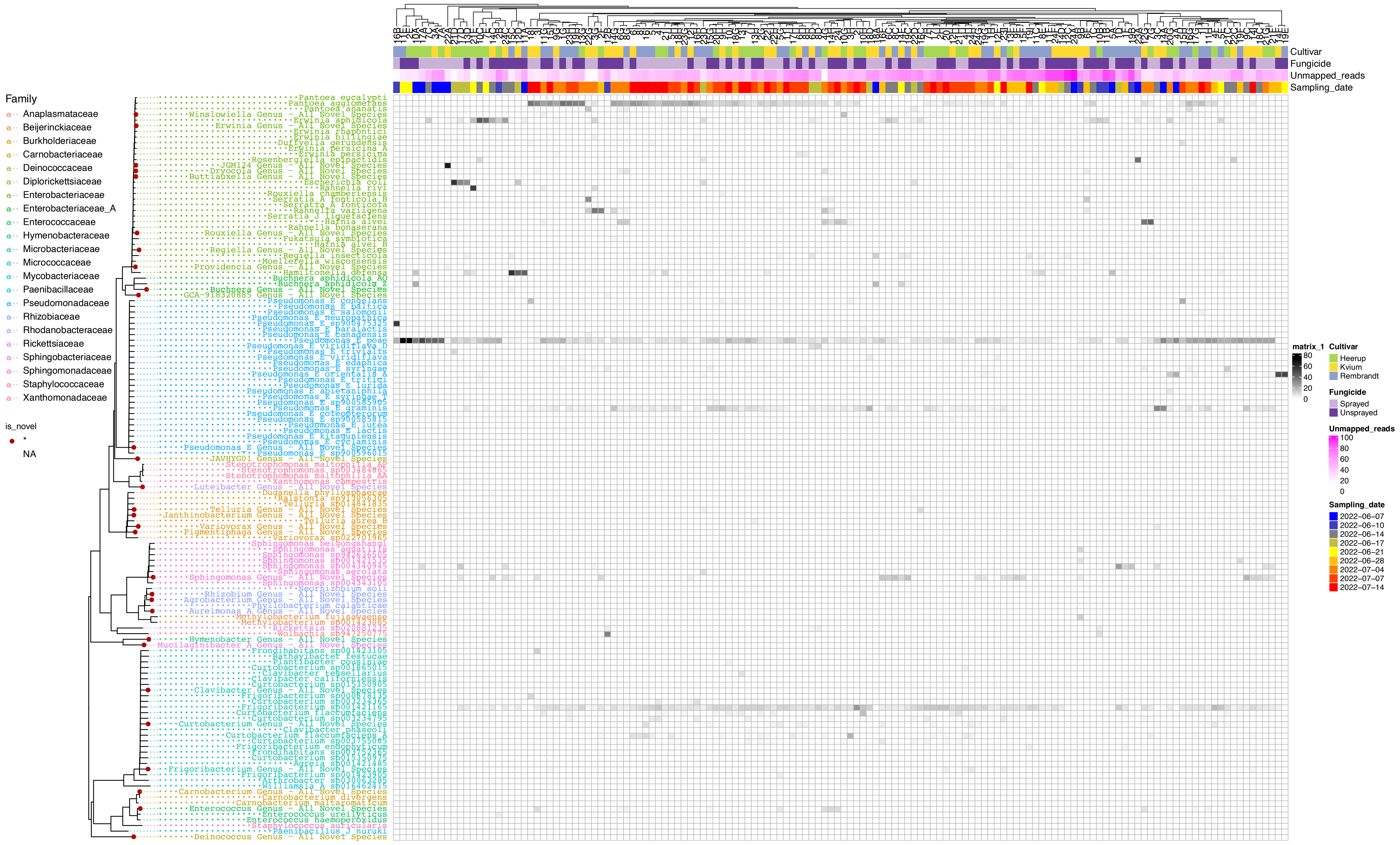


**Supplementary Figure S9 Abundance matrix for all species and samples, wheat phyllosphere.** Left, rows: species, organised in the phylogenetic tree reported by GTDB-tk. Top, columns: samples with high and medium quality MAGs, sorted by hierarchical clustering. The reported values are the shares of reads that were mapped to MAGs corresponsing to a species (sum for all the species MAGs for this sample) with completeness $\geq$ 50%, contamination $<$ 10% measured by CheckM2. On the top there are additional sample annotations: wheat cultivar, use of fungicide, share of unmapped reads, sampling date.

**Supplementary Table 1 Metagenome binners runtimes**. Evaluated in minutes, rounded down to integer, using the CAMI2 Oral dataset (10 samples)

| **Binner** | **Walltime (minutes)** | **Number of CPUs** |
|---|---:|---:|
| Metabat | 18 | 4 |
| MetaDecoder | 84 | 20 |
| SemiBin2 | 432 | 20 |
| AVAMB | 283 | 20 |
| VAMB (no reclustering) | 22 | 4 |
| TaxVAMB with Kraken2 labels (no reclustering) | 50 | 4 |
| Stacked VAE with Kraken2 labels (no reclustering) | 28 | 4 |
| Reclustering step (short-read) | 84 | 4 |

| Species (GTDB) | Prevalence | Association with plants from previous works |
|---|---|---|
| Pseudomonas E poae | 60% | antifungal activity, plant growth promotion [1] |
| Frigoribacterium sp001421165 | 53% | presence of Frigoribacterium genus positively correlates with wheat leaf size [2] |
| Pseudomonas E graminis | 41% | protective activity against leaf bacterial disease [3] |
| Pantoea agglomerans | 37% | antifungal activity, wheat growth promotion [4] |
| Erwinia aphidicola | 30% | pathogenic for aphids [5,6] |

**Supplementary Table 2 Wheat phyllosphere species prevalence.** Top 5 species sorted according to prevalence (share of samples with the species present with >1% of the mapped reads). Reads were mapped to MAGs with completeness $\geq$ 50%, contamination $<$ 10% measured by CheckM2. Columns: species name; prevalence (in % of the samples); brief descriptions of associations with phyllosphere as described in previous works.

| Genus (GTDB) | Prevalence |
|---|---|
| Pseudomonas E | 63% |
| Frigoribacterium | 54% |
| Pantoea | 39% |
| Erwinia | 37% |
| Curtobacterium | 28% |

**Supplementary Table 3 Wheat phyllosphere genus prevalence.** Top 5 genus sorted according to prevalence (share of samples with the species present with >1% of the mapped reads). Reads were mapped to MAGs with completeness $\geq$ 50%, contamination < 10% measured by CheckM2. Columns: genus name; prevalence (in % of the samples).
